# Imaging biomarkers for motor outcome after stroke – should we include information from beyond the primary motor system?

**DOI:** 10.1101/2020.07.20.212175

**Authors:** Christoph Sperber, Johannes Rennig, Hans-Otto Karnath

**Affiliations:** Centre of Neurology, Division of Neuropsychology, Hertie-Institute for Clinical Brain Research, University of Tübingen, Tübingen

## Abstract

Hemiparesis is a common consequence of stroke to the primary motor system. Previous studies suggested that damage to additional brain areas might play a causal role in the occurrence and severity of hemiparesis and its recovery. Knowledge of these regions might be applied in the creation of imaging biomarkers for motor outcome prediction if lesion information from such areas carries predictive value. We assessed acute and chronic paresis of the upper limb in 102 patients with unilateral stroke. In a first experiment, the neural correlates of acute and chronic upper limb paresis were mapped by lesion behaviour mapping. Following the same approach, a lesion biomarker of corticospinal tract (CST) damage was mapped. This analysis served as an artificial control condition as the biomarker, by definition, is only causally related to damage of the CST. Mapping acute or chronic upper limb paresis implicated areas outside of the primary motor system. Likewise, mapping the CST lesion biomarker implicated several areas outside of the CST with high correspondence to areas associated with upper limb paresis. Damage to areas outside of the primary motor system thus might, to some degree, not play a causal role in hemiparesis. In a second experiment, we showed that lesion information from these areas outside of the primary motor system can be used to predict motor outcome. This was even the case for the CST lesion biomarker. Although the only causal source underlying the CST lesion biomarker was damage to the CST, lesion information that mainly included non-CST regions was able to predict the biomarker (non-significantly) better than information taken from only the CST itself. These findings suggest that simple theory-based biomarkers or qualitative rules to infer post-stroke outcome from imaging data might perform sub-optimally, as they do not consider the complexity of lesion data. Instead, high-dimensional models with data-driven feature selection strategies might be required.

## Introduction

Brain lesions through strokes are a leading cause of acquired disability, and motor deficits such as weakness or paresis of the limbs are among its most common and debilitating symptoms. Such deficits arise after damage to the primary motor system, i.e. the primary motor cortex and its projection fibre, the corticospinal tract (CST). It is widely assumed that damage of the corticospinal tract underlies persisting primary motor deficits (Lindenberg *et al*., 2010; Lo *et al*., 2010; Zhu *et al*., 2010; Park *et al*., 2016). Thus, strategies to predict long-term motor outcome focused on assessing CST integrity by brain imaging (Lindenberg *et al*., 2010; Stinear *et al*., 2012; Wang *et al*., 2012; Sterr *et al*., 2014; Buch *et al*., 2016) or neurostimulation (Stinear *et al*., 2012).

Several studies developed prediction algorithms for motor outcome solely based on measures of CST damage derived from clinical CT or MR imaging (Wenzelburger *et al*., 2005; Zhu *et al*., 2010; Feng *et al*., 2015; Findlater *et al*., 2019). Metrics of CST lesion load were derived by referencing topographical lesion data to probabilistic topographies of the CST obtained by fibre tracking in healthy controls (Zhu *et al*., 2010; Feng *et al*., 2015; Findlater *et al*., 2019). These metrics were used in low-dimensional regression methods to predict motor outcome with moderate precision. However, two prediction studies using high-dimensional regression methods achieved better predictions if voxel-wise features were not only selected within the primary motor system, i.e. the CST and primary motor cortex, but, additionally, from a set of frontal, subcortical, and parietal areas (Rondina *et al*., 2016; Rondina *et al*., 2017). The authors hypothesized that the integrity of these regions plays a role in both motor deficits and motor recovery for their relation to motor and sensory function according to the literature. In other words, they performed a theory-driven feature selection, which resulted in improved prediction performance. A similar study found that in patients with stroke not confined to subcortical areas, using features taken from outside the CST improves model quality (Park *et al*., 2016). These findings complement lesion-behaviour mapping studies that found neural correlates of hemiparesis outside primary motor regions. Disintegrity of a cingulo-opercular attention control network was found to contribute to motor deficits (Rinne *et al*., 2017). Association fibres, the insula, and perisilvian areas were found to be associated with primary motor deficits in right hemisphere stroke, underlining a possible role of right hemisphere brain networks in motor recovery (Frenkel-Teledo *et al*., 2019). Another study found putamen and several fibre tracts to be associated with poor motor outcome (Findlater *et al*., 2019). Taken together, these findings suggest that stroke to regions outside the primary motor system might contribute to primary motor deficits. The identification of such regions could also guide researchers in creating imaging biomarkers for motor outcome prediction if imaging information taken from these areas bears predictive value.

A possible objection against the relevance of non-primary motor regions comes from critical discussions on lesion-deficit inference. The typical lesion anatomy was found to bias topographical results in lesion behaviour mapping (Mah *et al*., 2014; Sperber *et al*., 2019). It was postulated that the differentiation between areas where brain damage causes a deficit and areas where damage simply coincides with a deficit is impossible with current implementations of lesion-deficit inference (Sperber, 2020; but see Mah et al., 2014 for an opposing view). Thus, the causal interpretation of these regions’ role found in previous mapping studies might be– at least in parts – a misattribution. On the other hand, information outside of brain regions where damage causes a symptom can bear predictive value for a symptom (Sperber, 2020). Hence, in the face of this complexity, current biomarkers and feature selection strategies in motor outcome prediction might fall short if they are created in a theory-driven fashion, e.g. if motor outcome prediction is based only on information taken from regions where we assume damage to actually cause a long-lasting deficit.

The present study aimed to re-evaluate brain lesion-behaviour associations and prediction in upper limb paresis in 102 stroke patients. In the first experiment, we followed a pure mapping approach. We mapped the neural correlates of acute and chronic upper limb paresis. Additionally, we mapped a CST lesion load biomarker. The latter, per definition, is only causally related to damage to the CST. Therefore, this variable served as a perfectly understood reference condition, where unambiguous conclusions about mapping and prediction methodology, as well as causal relation in the data, could be drawn. We hypothesized upper limb paresis to be associated with areas outside the primary motor system, either because damage in these areas contributes in some way to primary motor deficits, or because damage coincides with damage to the primary motor system. Further, we also hypothesized CST lesion biomarkers to be associated with regions outside the CST, which would hint at brain areas where damage does not necessarily play any causal role for hemiparesis. In the second experiment, we evaluated how this affects prediction by using these areas found in experiment 1 as features in an out-of-sample prediction.

## Methods

### Patient recruitment and motor assessment

Patients with a first unilateral cerebral stroke admitted to the Centre of Neurology at the University of Tübingen were recruited. The diagnosis was confirmed by CT or MRI imaging. Patients with a medical history of other neurological or psychiatric disorders were excluded. All patients were re-examined in the chronic post-stroke stage at least 50 days and on average 423 days after the incident, and patients who suffered a second stroke were excluded. The final patient sample included 102 of which 68 had a stroke to the right, and 34 to the left hemisphere. The study has been performed according to the standards laid down in the revised Declaration of Helsinki and patients or their relatives consented to the scientific use of the data.

Paresis of the upper limb was assessed by the BMRC (British Medical Research Council) scale on average 1.0 days (SD = 2.2 days) after stroke onset. The BMRC scoring ranges from zero to five (0: no movement, 1: palpable flicker, 2: movement without gravity, 3: movement against gravity 4: movement against resistance, 5: normal movement) and allows intermediate decimals in steps of 0.5. It was used to assess paresis of the whole upper limb respectively distal and proximal parts. Ratings for distal and proximal upper limb motor function were averaged per patient. Demographic and clinical data of all patients are shown in Table 1.

**Table 1.**
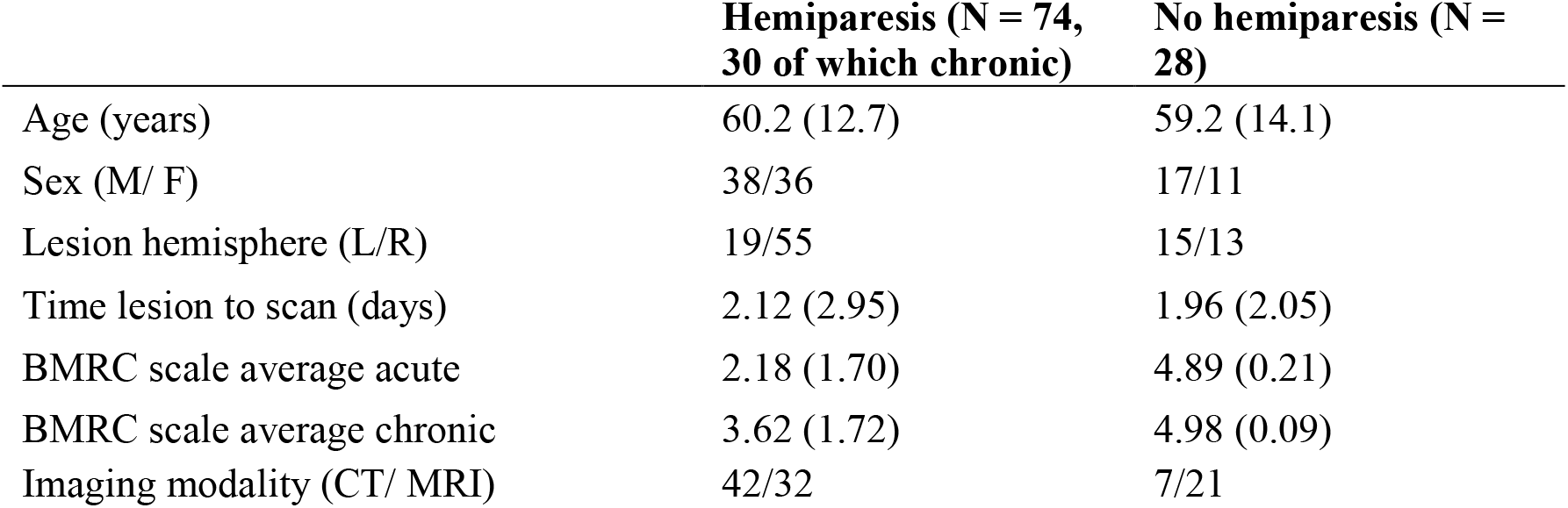
Demographic data. We defined a pathological range of the acute British Medical Research Council (BMRC) scale at scores <= 4. With this criterion, 74 patients were classified as hemiparetic, 30 of which presented chronic hemiparesis, and 28 had no motor deficit. Data are presented as mean (SD). Note that the binarisation of the data serves as a documentation for the inclusion of controls without motor deficits, while in the analysis, motor scores were treated as a continuous measure.

**Table 2.**
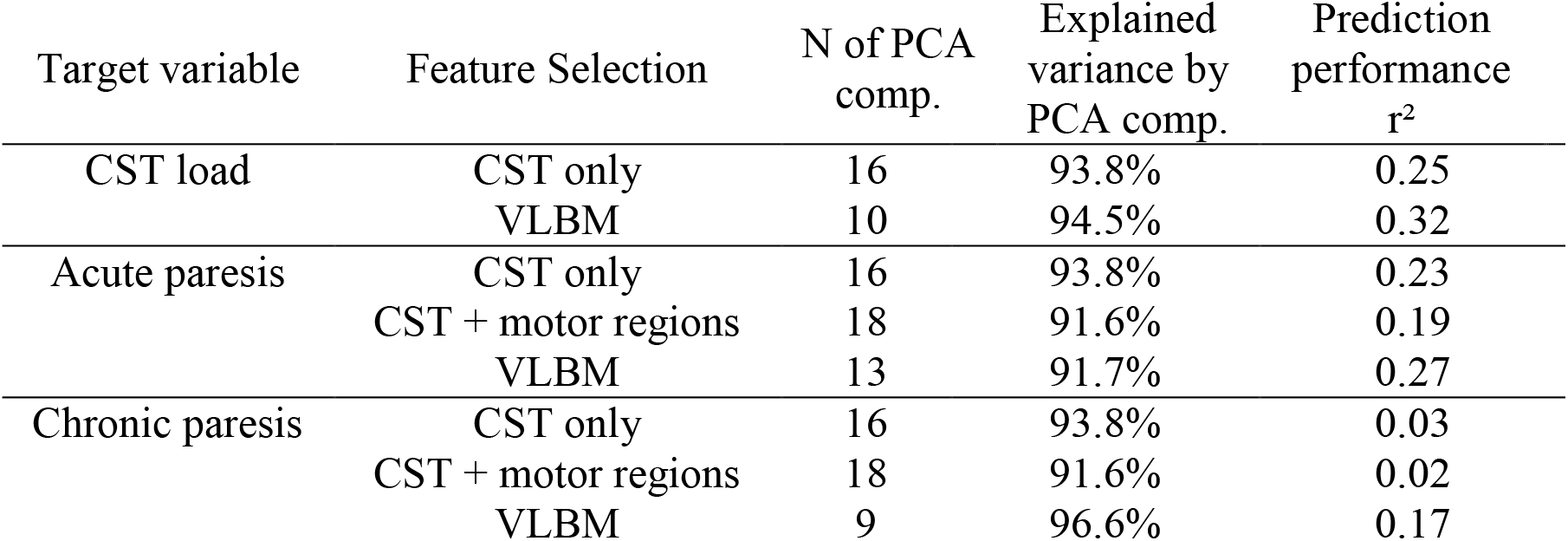
Results of experiment 2. Prediction performance was measured by the squared correlation between true scores and the scores predicted in the out of sample prediction. Explained variance by PCA components is the cumulative variance of explained lesion data by the retained components.

### Imaging and lesion delineation

Structural brain imaging was acquired by clinical CT or MRI on average 2.1 days (SD = 2.7 days) after stroke onset. If multiple scans were available, scans were chosen to be as acute as possible, while clearly depicting the borders of the lesion. If both suitable MRI and CT were available, MR scans were preferred. To visualize acute structural brain damage using MRI, diffusion-weighted imaging was used in the first 48h after stroke onset and T2 fluid-attenuated inversion recovery imaging afterwards. If available, these images were co-registered with a high-resolution T1 MRI to be used in the normalization process.

Lesions were manually delineated on axial slices of the scans using MRIcron (https://www.nitrc.org/projects/mricron) by two experimenters, and the accuracy of lesion drawings was validated by consensus with the third experimenter. Scans were warped into 1×1×1 mm^3^ MNI coordinates using Clinical Toolbox (Rorden *et al*., 2012), which contains age-specific CT and MRI templates in MNI space, and SPM 12 (www.fil.ion.ucl.ac.uk/spm), using cost-function masking to control for the lesioned area in the normalization process. The same normalization parameters were then applied to the lesion maps to obtain normalized binary lesion maps. Next, lesion maps of left hemisphere stroke patients were flipped along the sagittal mid-plane so that all lesions were depicted in the right hemisphere. This step was chosen as i) the study did not aim to investigate possible hemisphere-specific contributions to primary motor skills, but general methodological aspects in mapping primary motor functions and recovery, and as ii) this ensured high statistical power and adequate sample sizes for multivariate lesion-behaviour mapping. Figure 1 shows a topography of all normalized lesion maps.

**Figure 1:**
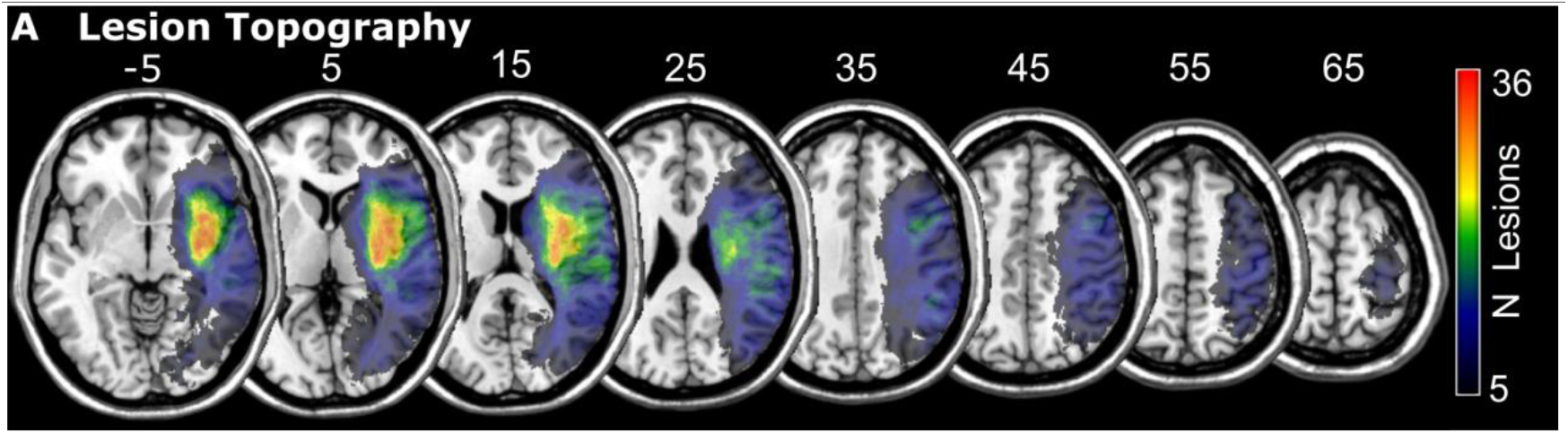
Lesion topography. Overlap topography of all 102 normalized binary lesion maps. The total number of overlapping lesions per voxel is depicted. Only voxels damaged in at least 5 patients are shown, which corresponds to the voxel included in the analyses. Numbers above the slices indicate z-coordinates in MNI space.

### CST lesion biomarkers

We followed previous studies that computed CST lesion metrics to obtain biomarkers for post-stroke motor outcome (Zhu *et al*. 2010; Feng *et al*., 2015; Findlater *et al*., 2019) and computed variables ‘CST load’ and ‘weighted CST load’. Both metrics assess to what degree a structural lesion affects the CST by computing overlap metrics between a normalized lesion map and a probabilistic topography of the CST. The required probabilistic topography of the CST was taken from a publicly available white matter atlas based on diffusion tensor imaging data in 81 healthy subjects (Mori *et al*., 2008). This topography indicates for each voxel in imaging space a probability between 0 and 1 to belong to the CST. For ‘CST load’ (Zhu *et al*., 2010), the overlap between lesion map and the probabilistic CST map is weighted by a voxel’s probability to be part of the CST. Accordingly, lesioned voxels that are more likely to be part of the CST contribute more to the CST lesion metric. For ‘weighted CST load’ (Zhu *et al*., 2010), the overlap between lesion map and the probabilistic CST map is weighted by the probability of a voxel to be part of the CST, and each voxel’s contribution is weighted by the cross-sectional area of the CST on each axial slice of the brain atlas. Accordingly, lesioned voxels in dense parts of the CST fibre bundle contribute more to the CST lesion metric. Computations were performed in MATLAB 2019b using SPM 12 algorithms. For more details on CST lesion metrics, see the formulas in Zhu *et al*. (2010) and the current study’s documented scripts, as available in the online materials. Final results for both CST biomarkers were highly similar, and thus results for weighted CST load are reported in the supplementary.

To ensure that topographical signal strength, as assessed by maximum Z values in the univariate lesion behaviour mapping analysis, is comparable between upper limb paresis (acute paresis max Z = 6.77; chronic paresis max Z = 6.58) and CST load (max Z = 8.25), random noise was added similar as in recent studies (Pustina *et al*., 2018). Noisy CST biomarkers were created by choosing a noise factor n in the range 0 <n < 1 and calculating noisy biomarkers nCST_bm_ based on original biomarkers CST_bm_ and uniform random values between 0 and 1 by 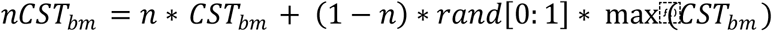. Factor n was tested in steps of 0.05 and manually chosen to provide optimal comparability between analyses of upper limb paresis and CST lesion biomarkers. For more information, see the online materials of the present study. The maximum Z value for noisy CST load (max Z = 6.92) was comparable in signal strength to upper limb paresis. Note that both original and noisy CST lesion biomarkers were mapped and interpreted.

### Data availability

Online materials including all analysis scripts, descriptive data, resulting topographies, multivariate lesion-behaviour mapping results, and full feature weight maps, are publicly available at http://dx.doi.org/10.17632/2pj8nxwbxr.2. The clinical datasets analysed in the current study are not publicly available due to the data protection agreement approved by the local ethics committee and signed by the participants. They are available from Hans-Otto Karnath on reasonable request.

## Experiment 1: Neural correlates of upper limb paresis and CST lesion biomarkers

### Methods - Lesion behaviour mapping

The association between voxel-wise lesion status and motor variables was mapped by univariate voxel-based lesion behaviour mapping (VLBM). The motor variables also included CST load, which is not a behavioural variable, but an imaging biomarker that was directly derived from the lesion image. Inclusion of this variable served as an artificial reference condition, closely resembling empirical validation strategies in lesion behaviour mapping (for review see Sperber & Karnath, 2018). We have perfect knowledge of the causality behind this variable, which allows us to unambiguously interpret results in the context of causality. All analyses were conducted at p = 0.05 only including voxels affected in at least five patients. VLBM was performed with voxel-wise general linear models, in the current study equivalent to t-tests, to investigate the relation between a voxel’s binary lesion status and the behavioural target variable in NiiStat (https://www.nitrc.org/projects/niistat/). Family-wise error correction for multiple comparisons was applied by maximum statistic permutation thresholding with 1.000 permutations. An additional multivariate analysis was also performed. Due to inferior performance, it is only reported in the online materials. Largest clusters in the topographical statistical results were assigned to brain regions by reference to the AAL atlas and a selection of fibre tracts found in the JHU white matter atlas in mricron (https://www.nitrc.org/projects/mricron), and the probabilistic map of the CST (Mori *et al*., 2008). For detailed descriptive data and underlying topographies, see the online materials.

## Results

Mapping the neural correlates of acute upper limb paresis with VLBM (Fig. 2A) found significant voxel centred on the CST, but also extending beyond the CST into insula, putamen, rolandic (fronto-parietal) operculum, postcentral gyrus, superior temporal gyrus, hippocampus, external capsule, corona radiata, and superior longitudinal fasciculus. Mapping chronic upper limb paresis with VLBM (Figure2B), no significant results were located in the CST. Instead, small clusters were found located in post- and precentral gyrus, and rolandic operculum.

**Figure 2:**
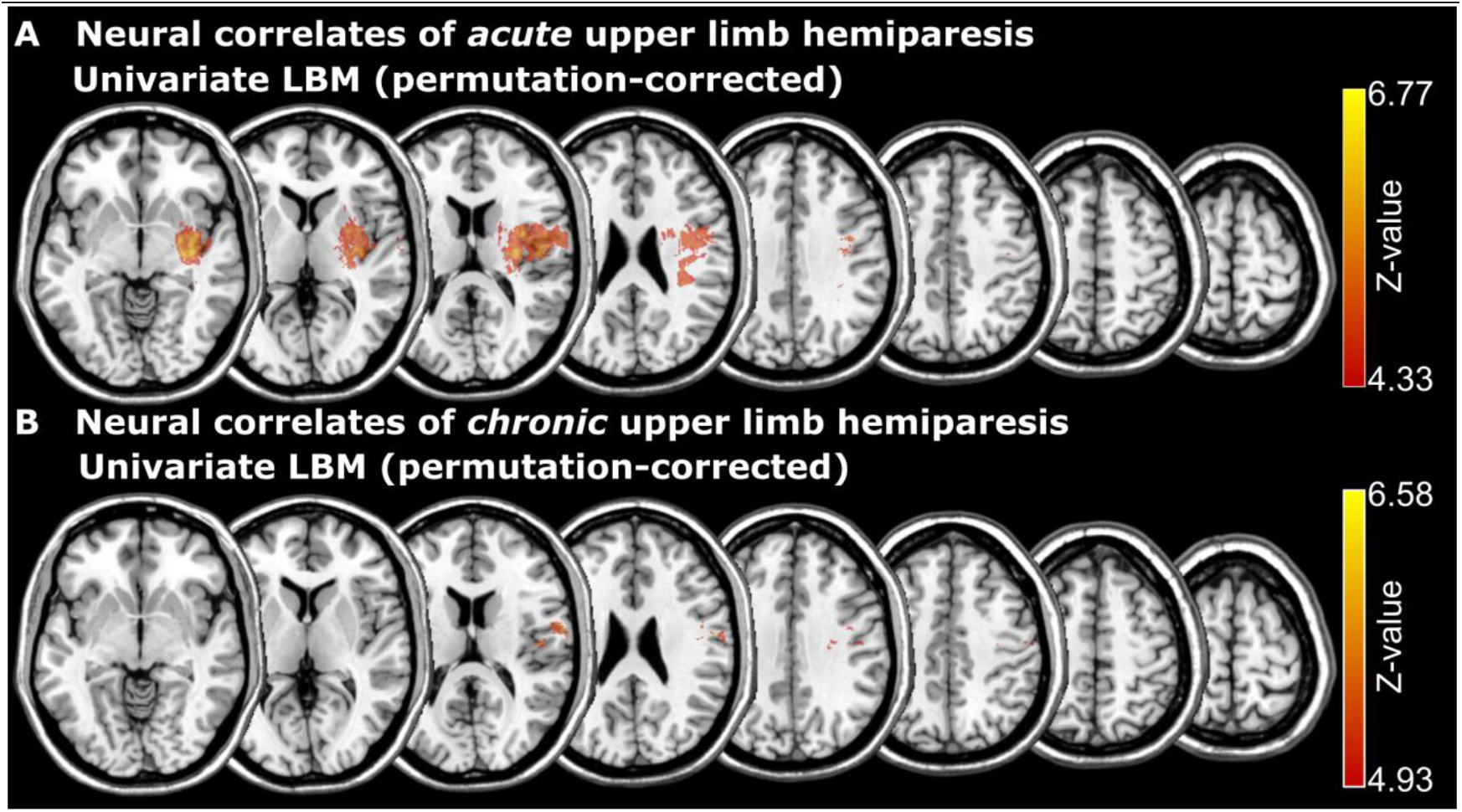
Lesion mapping of upper limb paresis. Voxel-wise lesion behaviour mapping of upper limb paresis both for A) acute and B) chronic motor assessment.

Mapping the neural correlates of ‘CST load’ without noise using VLBM (Figure3A) found significant cluster in the CST, but also extending into pre- and postcentral gyri, inferior, middle, and superior frontal gyri, supramarginal gyrus, rolandic operculum, superior temporal gyrus, putamen, pallidum, thalamus, superior longitudinal fasciculus, and corona radiata. Mapping the neural correlates of CST load with noise found, as expected, less significant voxels, due to reduced signal(Figure3B). Significant voxels were found in pre- and postcentral gyrus, rolandic operculum, putamen, supramarginal gyrus, external capsule, corona radiata, and superior longitudinal fascicle.

**Figure 3:**
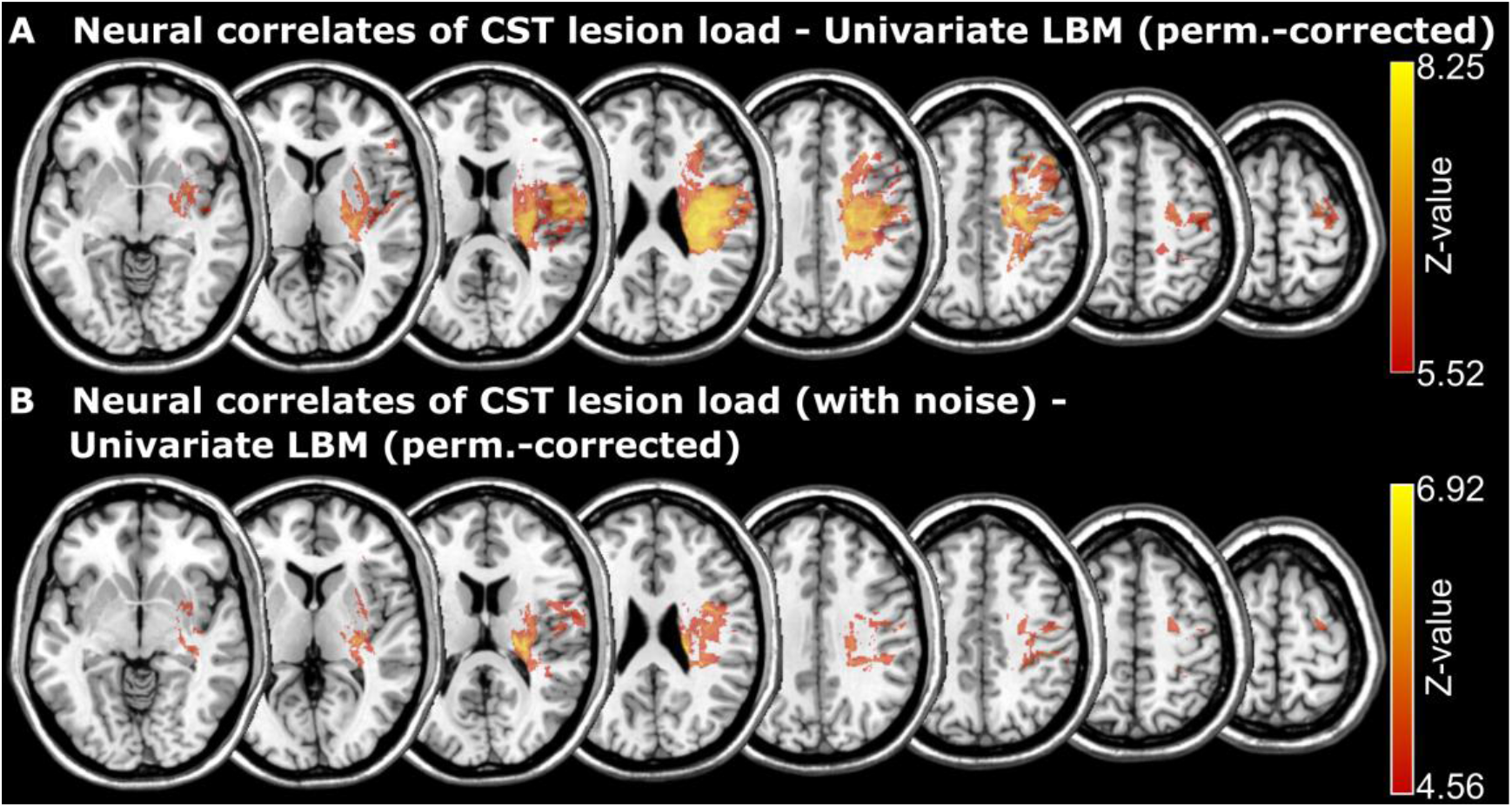
Lesion mapping of CST load. Voxel-wise lesion behaviour mapping of CST load both for A) raw lesion load scores and B) lesion load scores with noise added to match peak statistical signal with the mapping of upper limb paresis scores.

Results for weighted CST load (Supplementary Figure 1) closely resembled results for CST load and were thus not consulted in the data interpretation process. Topographies can be found in the supplementary and in the online materials of the present study. Next, VLBMtopographies of acute upper limb paresis and CST lesion biomarkers with noise were directly compared(Figure 4). This comparison was chosen as it mirrors a typical design to map the human brain (Karnath & Rennig, 2017), where univariate statistics are commonly applied on acute stroke data. Several areas outside the CST were found to be associated with both variables, including rolandic operculum, putamen, external capsule, superior longitudinal fascicle, and corona radiata.

**Figure 4:**
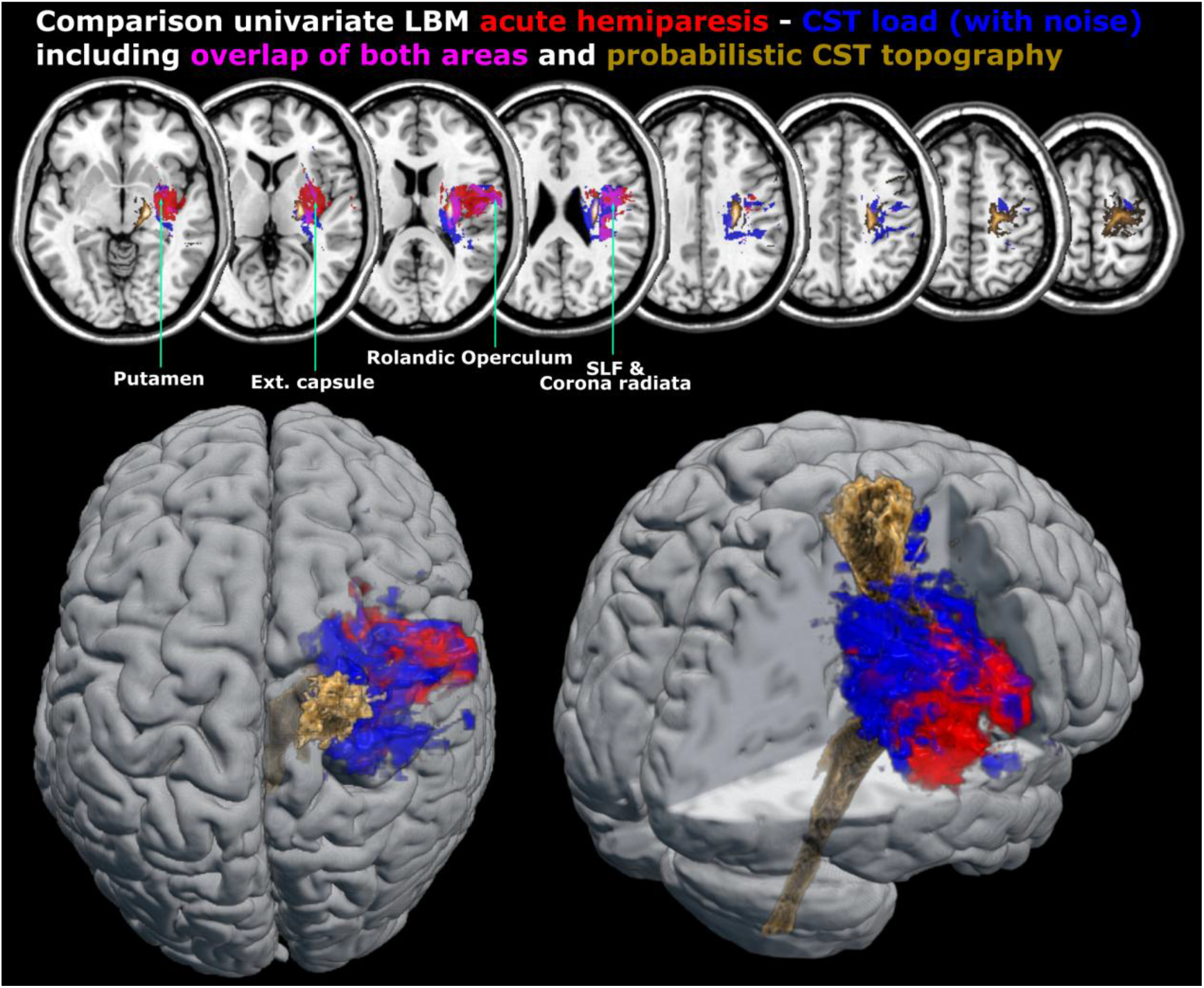
Comparison of the neural correlates of upper limb paresis and CST load. Direct comparison of statistical topographies for univariate VLBM between acute upper limb paresis and CST load (with noise). For reference, the probabilistic topography of the CSTis shown. Areas of overlap are indicated and labelled. SLF = superior longitudinal fascicle

## Discussion

We mapped the neural correlates of upper limb paresis and CST lesion biomarkers. The latter, per definition, only represent damage to the CST. Mapping acute or chronic upper limb paresis revealed areas outside of the primary motor system. This could suggest that upper limb paresis is not only caused by damage to the primary motor system, but also by damage to additional brain areas. However, mapping CST lesion biomarkers also implicated several areas outside of the CST, with some correspondence to the areas implicated in upper limb paresis. This hints at the possibility that non-primary motor brain regions can be found to be associated with hemiparesis not due to a direct causal contribution to the pathology, but due to lesion anatomy, or, more exactly, brain areas that are commonly damaged with primary motor areas. A possible clinical implication is that lesion information from these areas could meaningfully contribute to high-dimensional models for long-term motor outcome. In other words, damage to these areas might – at least to some extent –not play a causal role in hemiparesis, while still eventually providing predictive value for post-stroke motor outcome. To shed further light on this hypothesis, we conducted a second experiment, in which we predicted stroke outcome from lesion imaging data.

### Experiment 2: Modeloptimisation by feature selection

The second experiment aimed to clarify if outcome prediction with lesion information from outside the corticospinal tract and motor-related regions described in previous studies (Rondina *et al*., 2016; Rondina *et al*., 2017) can be improved by data-driven feature selection. Such feature selection poses a complex challenge. Therefore, the current study aimed to only provide a proof of principle by using the topographical information obtained in experiment 1 to select features. This is not an optimal solution, as features that are found to be significantly associated with a target variable and features that are predictive are not equivalent (Bzdok et al., 2020). Nonetheless, positive results already with such a suboptimal approach would make a clear point for data-driven feature selection. Three different variables were predicted from lesion imaging: acute hemiparesis, chronic hemiparesis, and CST lesion load. Again, the variable CST lesion load served as an artificial reference condition without an actual behavioural measure.

## Methods – Feature selection

Features were extracted on a voxel-wise level from normalised lesion imaging data. For each patient, imaging features were extracted from i) the CST, ii) the CST and motor regions as described by Rondina and colleagues (2016), or iii) from voxels found to be associated with the target variable by univariate voxel-based lesion behaviour mapping. The CST was again taken from a DTI atlas (Mori *et al*., 2008), and a binary map was created from the probabilistic map by including all voxels with a probability to be part of the CST > 0. A previous study found that features taken from a set of motor-related regions were better suited in prediction than only features taken from the CST alone (Rondina *et al*., 2016). These regions were defined with the Automated Anatomical Labelling Atlas (AAL; Tzourio-Mazoyer *et al*., 2002) and included postcentral gyrus, precentral gyrus, supplementary motor area, superior frontal gyrus, middle frontal gyrus, Inferior and Superior parietal regions, Thalamus, Caudate, Putamen and Pallidum (see Figure 5A). This feature selection was only used to predict acute and chronic hemiparesis. Voxels associated with a target symptom in univariate VLBM were defined by the corresponding maps from experiment 1 after correction for multiple comparisons (see Figure 3A) provided widespread clusters and large effects, which far surpassed the signal strength in the comparable analyses on actual behavioural scores. Features taken from this map were found to perform sub-optimally, in accordance with the partial discrepancy between brain mapping and prediction (Bzdok *et al*., 2020). We manually tested more conservative statistical thresholds and found a cut-off at a Z-value>7.3 (Figure 5B for a topography) to provide features for better model performance. This topography partially overlapped with the CST map, however, the majority of voxels (74.8%) were not part of the CST map.

**Figure 5:**
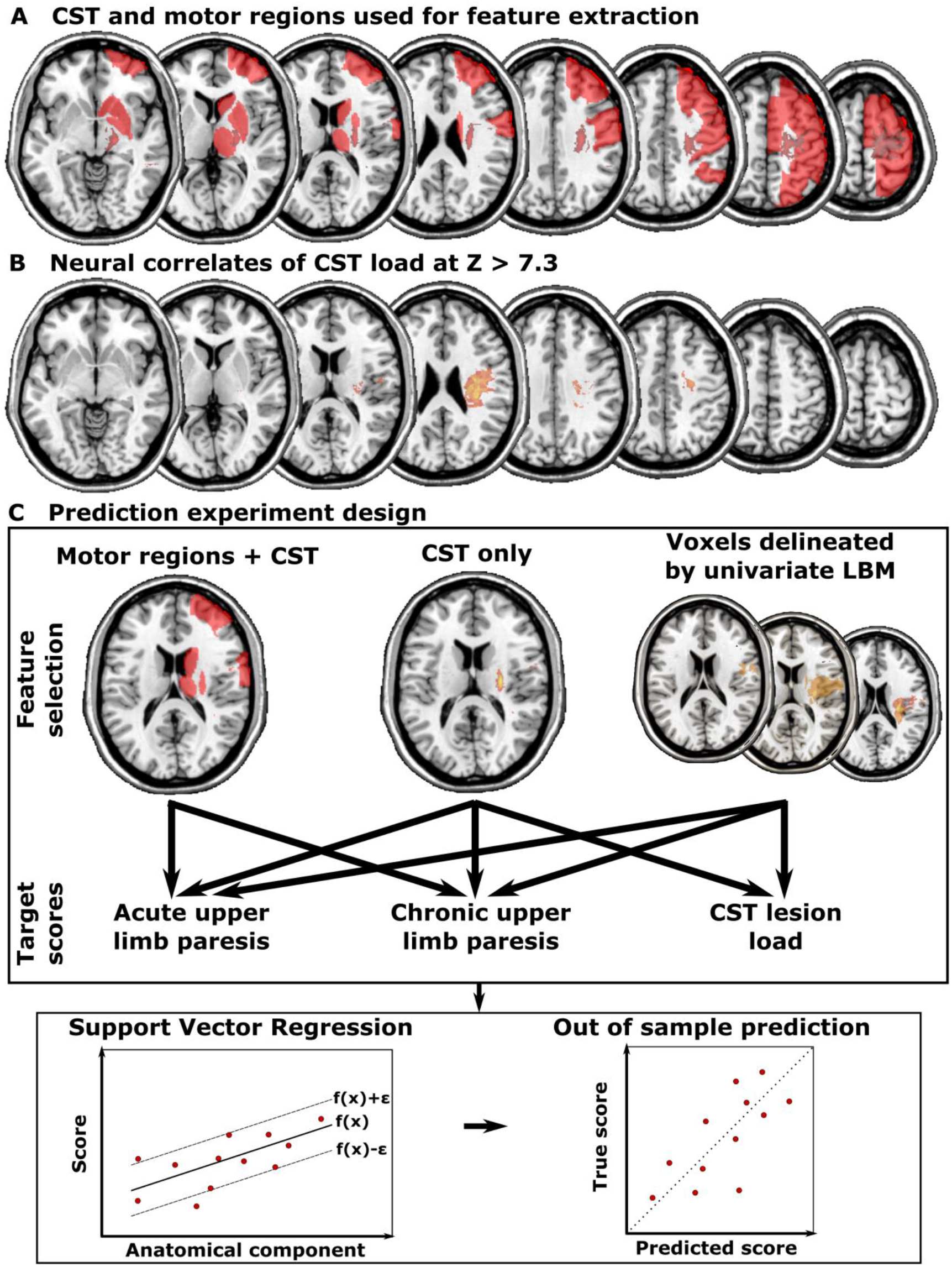
Extracted features and design of experiment 2. A) Motor regions and CST. B) Features delineated by the univariate LBM of CST load at the adapted threshold at Z > 7.3. C) Design of experiment 2.

## Methods – post-stroke outcome prediction

After feature selection, the imaging data were subjected to principal component analysis. Only components explaining at least 1% of the total variance were retained. The remaining features were used in anε-support vector regression (SVR) to predict the three different target variables in libSVM 3.24 (Chang and Lin, 2011). With the default ε = 0.1 (as used for lesion data in Zhang *et al*., 2014), we optimised hyperparameters C and γ in the range of C = [2^-3^, 2^-2^, …,2^10^] and γ = [2^-10^, 2^-9^, …, 2^3^] in a grid search. Model performance was evaluated by computing the correlation r between predicted and true scores. The full dataset was divided into six folds with 17 subjects each. Hyperparameters were optimised within five out the six folds by computing an SVR with each pair of hyperparameters on four folds, and then model performance was evaluated in the fifth fold. This was done for all five folds, and overall hyperparameter performance was then averaged across all five folds. The pair of hyperparameters with the best performance was then used to again train a model on all five folds and predict the outcome in the remaining sixth fold. Following the same strategy, predictions were generated six times, resulting in a predicted score for each patient. Observations included in folds were the same across all analyses. Differences in out-of-sample prediction between feature selection strategies were evaluated for statistical significance by comparing residuals with the Wilcoxon rank test.

## Results

Across all analyses, between 9 and 18 components remained after principal component analysis. Cumulatively, components explained > 91% of the variance in all analyses. For all three target variables, voxels delineated by univariate VLBM provided the model with the best model performance (see Tab. 2). However, differences failed to reach statistical significance, and thus we only descriptively report differences. CST lesion load was indeed minimally better predicted by voxels delineated by VLBM (r^2^ = 0.32) than by voxels inside the CST (r^2^ = 0.25). Most notably, the prediction of chronic upper limb paresis profited from predictions based on voxels delineated by VLBM (r^2^ = 0.17) compared to voxels inside the CST (r^2^ = 0.03). This was especially surprising as the VLBM map did not include the CST at all. Features extracted from the CST and motor regions provided the lowest performance in the current study.

## Discussion

The selection of features by univariate voxel-based lesion behaviour mapping using general linear models cannot be seen as an optimal feature selection strategy (see Bzdok *et al*., 2020). Still, the current study provides a proof of principle that prediction following such data-driven strategy (non-significantly) outperforms theory-driven feature selection. Importantly, this was even the case for the purely methodological control condition where we mapped CST lesion load – a lesion imaging biomarker that only represents CST damage. CST load was better predicted using voxels found by the preceding VLBM. These voxels included some parts of the CST, but the majority of voxels were located outside the CST.

### General discussion

We investigated brain lesion-behaviour associations and prediction of post-stroke motor outcome, as well as a purely methodological control condition, in which we mapped a CST lesion load biomarker. This biomarker, per definition, only represents damage to the CST. Mapping acute or chronic upper limb paresis, areas outside of the primary motor system were revealed. However, mapping the association between brain lesions and CST lesion biomarkers also implicated several areas outside of the CST, with some correspondence to the areas implicated in upper limb paresis. Damage to these areas might not play a causal role in hemiparesis. The brain regions found in the mapping analysis also carried predictive value for the motor variables in an out of sample prediction. These predictions (non-significantly) surpassed the prediction performance when using information from only the CST or the CST and motor regions. Notably, this was even the case for the CST lesion biomarker, implying that there is some discrepancy between causality in brain lesion-outcome relations and predictive value. These findings impose the need to rethink feature selection strategies and biomarkers in post-stroke outcome prediction based on imaging data.

At first, it might appear counter-intuitive that CST lesion biomarkers are related to damage outside of the CST. This finding is based on the systematic complexity of brain lesions. Stroke follows the vasculature, resulting in typical stroke patterns, and neighbouring brain regions are systematically damaged together. Thus, even if upper limb paresis was caused only by damage to primary motor regions, mapping hemiparesis would implicate areas outside of the primary motor system. This caveat of all current lesion-deficit inference methods has been acknowledged recently (Mah *et al*., 2014; Sperber *et al*., 2019; Sperber 2020). Notably, experiment 2 showed that areas outside of the primary motor system are predictive of hemiparesis in high-dimensional models, even if they might not be directly causally related to the pathology. In general, predictions can be made based on variables that are not the direct cause of the target variable: we can predict the weather from a swallow’s flight, even though it is not the swallow that causes rain. The general need to differentiate between direct causality and predictive value in the feature selection of imaging biomarkers is illustrated with a fictional example in Figure 6.

**Figure 6:**
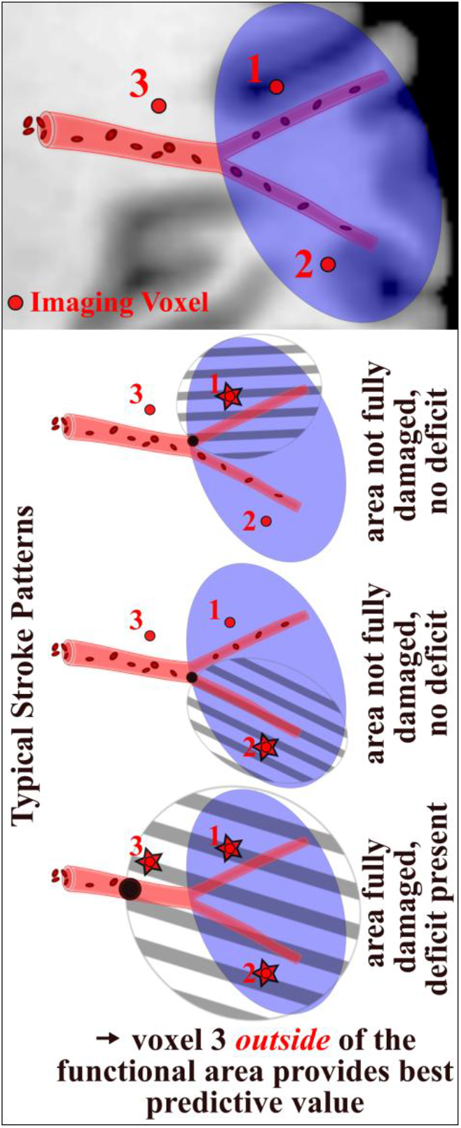
A fictional example to illustrate the discrepancy between functional brain anatomy and predictive value in stroke imaging biomarkers. The neural correlate of a behaviour (blue area) is supplied by two branches of an artery. Only full damage to the area evokes a deficit. Given the typical anatomy of lesions – after ischemia in either of the branches or the preceding artery – imaging information taken from inside the neural correlates (voxel 1 & 2) provide only sub-optimally predictive value for the deficit. On the other hand, information that is taken from outside the neural correlate (voxel 3) might even surpass the predictive value of single imaging features obtained from the area where damage actually caused the symptom.

Several previous studies suggested that features taken from regions outside of the primary motor system should be included in post-stroke outcome prediction (Park *et al*., 2016; Rondina *et al*., 2016; Rondina *et al*., 2017). The rationale behind this approach was theories that causally link these regions and primary motor deficits, e.g. the assumption that these areas are relevant for motor and sensory function (Rondina *et al*., 2016; Rondina *et al*., 2017). Indeed, prediction performance was improved by this strategy in previous studies, but the present study suggests that this might have happened for other reasons. Some of the proposed motor regions overlapped with regions that were found to be associated with CST biomarkers in the present study, including subcortical and frontal regions. Notably, in the current study, features taken from these motor regions were (non-significantly) inferior to features taken from the lesion-behaviour maps when used in outcome prediction. Several of the proposed motor areas were not found to be associated with motor deficits in the present study, and thus only a subset of regions might underlay possible improvements in prediction. The question remains which areas outside of the primary motor system play a direct causal role in primary motor deficits and if this was the case for at least some of the regions chosen by previous prediction studies. The current study was not designed to identify what areas constitute the true neural correlates of upper limb paresis. Our findings rather imply that topographical results have to be interpreted with caution, and favour the view that the combination of lesion-deficit inference with strong hypotheses and behavioural experiments (e.g., Rinne *et al*., 2017), or other imaging and neurostimulation techniques are required to provide a solid foundation to understand the architecture of the human motor system and the human brain in general (Rorden & Karnath, 2004).

Clinical imaging is part of any diagnostic protocol, and prediction algorithms using only such data would be widely applicable and economic, providing the potential to revolutionize individual patient care. However, no effective algorithms exist yet (Nachev *et al*., 2019). Low-dimensional prediction algorithms, such as simple linear regression, achieve up to moderate prediction performance at best based on CST lesion biomarkers (Findlater *et al*., 2019). For clinically useful algorithms, prediction performance has to be further improved, and high-dimensional models will likely be required to achieve this (Nachev *et al*., 2019). Feature selection is a vital step in any such algorithm based on machine learning. The current study indicates that this is a difficult task in stroke imaging data. High-dimensional datasets of massive size that contain highly correlated features simply cannot be precisely described by simple rules. Confident knowledge about causal relations between variables is even more difficult to obtain (Shiffrin, 2016). Brain imaging in stroke provides a prime example for such datasets. It appears that the brain and its functional organisation cannot be properly represented by single anatomical structures, and, correspondingly, that brain pathology cannot be understood without considering the brain’s complex network structure (Catani *et al*., 2012). This also implies that stroke outcome prediction might be further improved not only by better accounting for the complexity of functional brain anatomy on a voxel-wise level, as done in the present study, but also by representing the brain’s functional network structure (e.g., Griffis *et al*., 2019; Salvalaggio *et al*., 2020) and inter-regional functional relations (e.g., Toba *et al*., 2020). Brain pathology such as stroke adds another layer of complexity, as the pathological status between features is also highly correlated (Pustina *et al*., 2018; Sperber, 2020). Further, post-stroke brain plasticity introduces a longitudinal aspect to this complexity. Structural alterations in several grey and white matter regions were found to be associated with the relearning of motor skills (Sampaio-Baptista *et al*., 2018), which could also underlie a possible role of non-primary motor areas in post-stroke motor outcome.

## Conclusions

Selecting imaging features to predict post-stroke outcome based on our understanding of the brain might perform sub-optimally, even for – compared to the complex language or spatial attention systems – relatively well-understood and more easily accessible brain functions such as primary motor abilities. Instead, data-driven feature selection strategies seem to be required to improve prediction algorithms and bring us closer to effective predictive personalized medicine in stroke rehabilitation. The approach to data-driven feature selection in the present study only served as a proof of principle, and the optimisation of feature selection poses a future challenge for the field.

## Declaration of Competing Interests

The authors report no competing interests.

## Funding

This work was supported by the German Research Foundation (KA 1258/23-1).

## Notes

http://dx.doi.org/10.17632/2pj8nxwbxr.2

### Competing Interest Statement

The authors have declared no competing interest.

### Summary of Updates

We added a second experiment with out-of-sample prediction, making a much stronger point for the mansucript's implications for prediction. The previous multivariate mapping analysis was removed from the main manuscripts, but still can be found in the online materials. Further, several changes were made to improve clarity and readability.

