## Supplemental Figure for "Imaging biomarkers for motor outcome after stroke – should we include information from beyond the primary motor system?"

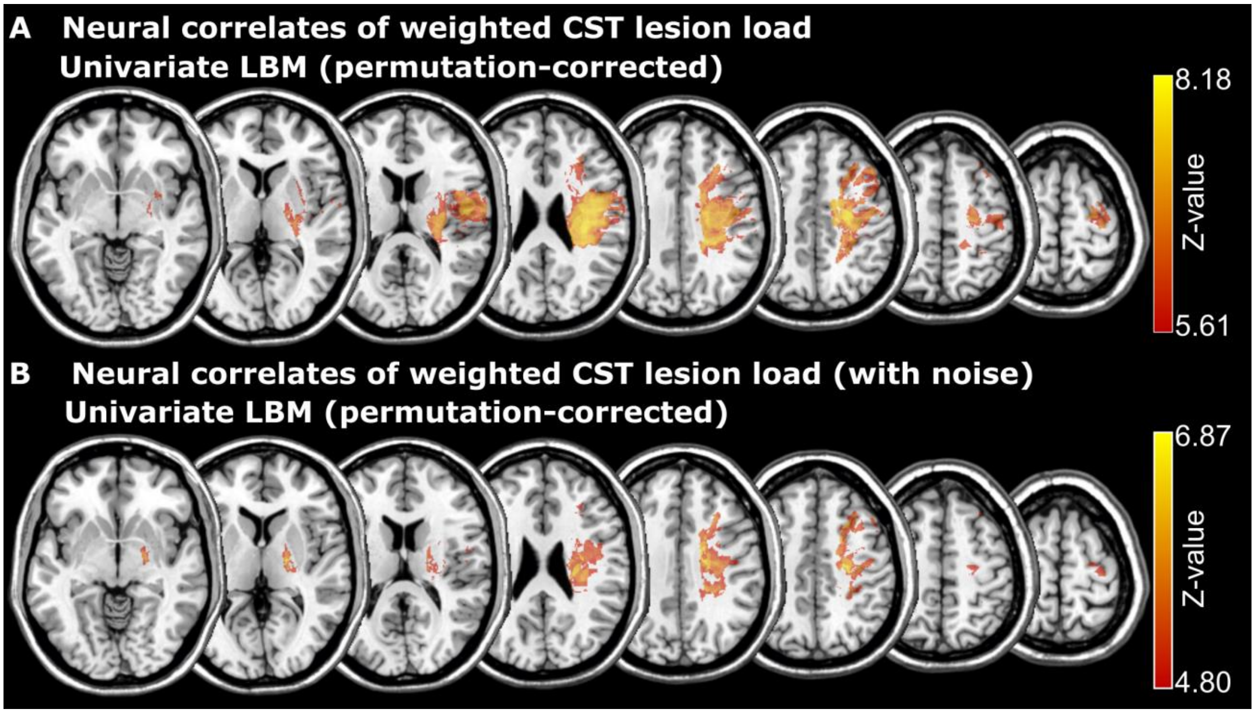


**Supplementary Figure: Lesion mapping of weighted CST load**

Voxel-wise lesion behavior mapping of CST load both for A) raw lesion load scores and B) lesion load scores with noise added to match peak statistical signal with the mapping of upper limb paresis scores. The maximum Z-values were max Z = 8.18 in the original map and Z = 6.87 after addition of noise.
